# Characterisation of distinct senescent-like T cell populations during healthy and unhealthy ageing

**DOI:** 10.1101/2025.06.16.659752

**Authors:** Conor Garrod-Ketchley, Lauren A Callender, Johannes Schroth, Katie Littlewood, Elizabeth C Carroll, Dina Tamsan, Victoria SK Tsang, Isabell Nessel, Natalie E Riddell, Benny Chain, Dunja Aksentijevic, Daniel Harding, Federica M Marelli-Berg, Jonas Bystrom, Melissa Pereira Da Costa, Amaia Carrascal-Miniño, George P Keeling, Truc T Pham, Kavitha Sunassee, Rafael TM de Rosales, Samantha YA Terry, Melanie Pattrick, Caroline Sutcliffe, Anne Worthington, Gill Hood, Sarah Finer, Sian M Henson

## Abstract

Ageing is accompanied by progressive remodelling of the immune system, but chronic metabolic disease may accelerate this process and drive qualitatively distinct forms of immune dysfunction. Here, we used type 2 diabetes (T2D) as a model of unhealthy immune ageing to identify a distinct population of CD8^+^ T_EMRA_ cells that accumulates in older individuals with T2D. These cells were highly differentiated, oligoclonally expanded and had shorter telomeres, consistent with an increased replicative history and premature senescence-like state. Unlike conventional T_EMRA_ cells, which can preserve cytotoxic function through acquisition of NK-cell receptors, T2D-associated T_EMRA_ cells showed reduced surface expression of KLRG1, NKG2D and NKG2A together with defective receptor recycling. TGFβ1 was elevated in T2D and reproduced several features of this phenotype in vitro, including increased T_EMRA_ differentiation, reduced NK-receptor expression, altered receptor trafficking and induction of p21. Functionally, these cells displayed reduced TCR-triggered degranulation, correlating with impaired cytotoxicity and altered tissue distribution in individuals with T2D. Rather than providing effective immune surveillance, the accumulation of these highly differentiated T_EMRA_ cells with diminished effector capacity may compromise immune function. Together, these findings identify a distinct senescent-like CD8^+^ T_EMRA_ state linking metabolic inflammation to dysregulated T cell differentiation.

## Introduction

For the first time in history, older individuals constitute the fastest growing demographic worldwide. Projections indicate that between 2015 and 2050, the proportion of the global population aged 60 years and older will nearly double, increasing from 12% to 22% (1). This prolonged life expectancy, together with the increased prevalence of age-related diseases, underscores the critical importance of maintaining an effective immune system throughout ageing.

A well-documented consequence of ageing is its profound impact on the T cell compartment (2-5). Due to their highly proliferative nature, CD8^+^ T cells are particularly vulnerable to the effects of ageing and show a high degree of compartmental heterogeneity. One notable consequence is the accumulation of naïve-like memory cells, virtual memory cells (T_VM_), which despite being antigen-inexperienced can exert effector functions (6, 7). Alongside this is the differentiation of the memory pool that leads to a population of CD8^+^ T cell that progressively loses functionality with age. The identification of T cell heterogeneity has relied heavily on phenotypic marker profiles, distinguishing stem cell memory cells (T_SCM_), central memory (T_CM_) and effector memory (T_EM_) cells along with terminally differentiated effector memory T cells that re-express CD45RA (T_EMRA_). This classification relies on the combinatorial expression of markers such as CD27, CD28, CCR7, CD45RA and CXCR3 (8). Senescent-like T_EMRA_ cells, in particular adopt innate-like properties, acquiring the expression of numerous NK receptors that enable TCR-independent cytotoxicity (9). The increased complexity of T cell populations with age has been further elucidated through advanced ‘omics’ technologies, such as single-cell cytometric and RNA-seq analysis. These approaches, like IMM-AGE and iAge, used in combination with phenotypic markers can model immune senescence providing a high-dimensional trajectory of immune ageing (10, 11).

The accumulation of T_EMRA_ cells is often driven by repeated antigen stimulation, particularly from chronic infections such as cytomegalovirus (CMV), resulting in memory T_EMRA_ cells with a senescence-associated phenotype (12, 13). Immune senescence represents a progressively degenerative state characterised by the loss of replicative capacity or onset of replicative senescence (2). This process is initiated by a DNA damage response (DDR) resulting from telomere shortening (14). Critically short telomeres activate the ataxia-telangiectasia mutated (ATM) and ataxia-telangiectasia and Rad3-related (ATR) kinases, leading to the phosphorylation of histone H2AX (γH2AX) and activation of downstream effectors such as p53, p21^CIP1/WAF1^ and retinoblastoma protein (Rb), which collectively enforce a stable cell-cycle arrest characteristic of replicative senescence (15). Beyond telomere erosion, other intrinsic and extrinsic stressors can also drive cellular senescence through stress-induced premature senescence (SIPS). Oxidative stress, DNA-damaging agents and oncogene activation can induce persistent DDR signalling independent of telomere attrition via activation of p16^INK4A^ and p53-p21 pathways (16). While SIPS is not directly driven by telomere shortening, persistent DNA damage signalling, particularly involving γH2AX and 53BP1 foci, can contribute to telomere dysfunction, potentially linking SIPS to telomere-dependent mechanisms in certain contexts (17).

Additionally, metabolic stability plays a key role in determining susceptibility to senescence. Disruptions in metabolic homeostasis through pathways that regulate energy balance, oxidative stress, and mitochondrial integrity all accelerate the onset of senescence (18). The activation of the metabolic sensor AMPK in senescent T cells, through DNA damage or reduced levels of ATP, heightens endogenous p38 phosphorylation which controls the inflammatory secretome associated with senescence (19). Furthermore, the decline in mitochondrial fitness observed in CD8^+^ T_EMRA_ cells increases dependency on glycolysis and elevates ROS production (20). These dysfunctional mitochondria lead to impaired energy production and the accumulation of oxidative stress, promoting senescence-associated phenotypes. Here, we distinguish biological ageing, defined as systemic physiological decline, from immune ageing, where metabolic dysregulation and low-grade inflammation may amplify ageing-related immune phenotypes.

Given that senescence is sensitive to both inflammatory and metabolic stress, we investigated the phenotype and function of T cells in people living with type 2 diabetes (T2D), a condition associated with premature ageing (21). T2D is characterised by a chronic metabolic and inflammatory environment that may accelerate or reshape aspects of immune ageing. Consistent with this, numerous studies have identified immune features in T2D that are also associated with ageing, including reduced naïve T cell output (22), expansion of senescent (23, 24) and exhausted T cell subsets (25), persistent low-grade inflammation (26) and impaired antigen-specific immune responses (27). Studies of adaptive immunity in diabetes further support substantial remodelling of T cell subset composition in association with systemic inflammation (28). Importantly, these changes should not be interpreted as evidence that T2D is equivalent to biological ageing. Rather, T2D provides a clinically relevant setting in which chronic metabolic dysfunction and inflammation may promote an accelerated or dysregulated immune-ageing phenotype. Indeed, previous analysis of CD8+ T cells in T2D identified features of accelerated immune ageing that were not fully explained by glycaemic control alone (29). Here we use T2D as a clinical model of unhealthy immune ageing, defined as a disease-associated trajectory shaped by persistent metabolic dysregulation, chronic inflammation and impaired immune homeostasis. This framework distinguishes unhealthy immune ageing from chronological or biological ageing and does not imply that T2D is representative of all forms of unhealthy ageing. Selected phenotypic observations were also examined in individuals with dilated cardiomyopathy to assess whether aspects of the phenotype extended beyond T2D.

Healthy ageing is generally associated with gradual remodelling of the immune system while retaining sufficient homeostatic control and immune surveillance (30). In contrast, unhealthy immune ageing may involve persistent low-grade inflammation, metabolic stress and immune dysregulation, promoting the accumulation of highly differentiated and senescent-like immune populations and contributing to declining immune competence. In this study, we compare healthy ageing with T2D-associated unhealthy immune ageing and identify a population of CD8^+^ T cells with features of premature senescence and altered differentiation. Functionally, T_EMRA_ cells and broader CD8^+^ T_EMRA_ cell populations from the T2D cohort exhibited altered tissue-associated characteristics and reduced responses following anti-CD3 stimulation. Together, these findings suggest that chronic metabolic and inflammatory stress is associated with a distinct trajectory of CD8^+^ T cell ageing and may contribute to impaired immune function during unhealthy ageing.

## Methods

### Ethics and donor recruitment

Approval for our study was granted by the West London & GTAC Research Ethics Committee (20/PR/0921) and all methods were carried out in accordance with approved guidelines and regulations. Written informed consent was obtained for all participants. We recruited three separate cohorts, healthy old participants and people living with T2D, identified through the Diabetes Alliance for Research in England (DARE) database (cohort 1). A cohort of individuals with dilated cardiomyopathy (DCM) with and without T2D (cohort 2) and individuals undergoing carotid endarterectomy (cohort 3) both of which were collected under ethics granted to the Barts Cardiovascular Registry (REC 14/EE/0007). See table 1 for full patient characteristics. Peripheral blood was obtained using heparinised tubes and peripheral blood mononuclear cells (PBMCs) were isolated using Ficoll hypaque (Amersham Biosciences), with frozen material stored in liquid nitrogen in a suspension of 10% DMSO to 90% FBS.

**Table 1.** Demographic and clinical characteristics of the study cohorts.

|  | Cohort 1 |  | Cohort 2 |  | Cohort 3 |
| --- | --- | --- | --- | --- | --- |
|  | Healthy | T2D | DCM | DCM + T2D | Carotid endarterectomy |
| Total number of participants | 120 | 120 | 36 | 16 | 4 |
| Age range (years) | 55-80 | 52-78 | 52-79 | 55-74 | 50-66 |
| Average age (years) | 70 ($\pm 7$) | 68 ($\pm 5$) | 65 ($\pm 7$) | 62 ($\pm 9$) | 58 ($\pm 6$) |
| Gender (male) | 49% | 56% | 77% | 68% | 75% |

### Flow cytometric analysis

Flow cytometric analysis was performed using the following antibodies: anti-CD8 PerCP (SK1), anti-CD45RA BV605 (HI100), anti-CD45RA APC (HI1000), anti-CD27 BV421 (O323), anti-CD27 FITC (O323), anti-CD28 BV785 (CD28.2), anti-CCR7 PECy7 (G043H7) anti-CD57 AF700 (HNK-1), anti-NKG2A AF700 (S19004C), anti-NKG2D (1D11), anti-CD49d PE (9F10), anti-CD103 APC (Ber-ACT8), anti-CD69 PE (FN50), anti-perforin APC (dG9), anti-granzyme B FITC (QA16A02) and anti-CD107a APC (H4A3) from BioLegend; anti-KLRG1 PE (MAFA) from Miltenyi Biotec; and anti-NKG2C AF488 (134591) R&D Systems. Live/Dead Fixable Near-IR 775 stain (ThermoFisher). Intracellular staining was carried out using solution AB (ThermoFisher).

All samples were analysed using a LSR Fortessa (BD Biosciences) and the resulting data examined using FlowJo software (BD Bioscience).

### Telomere length assessment by flow-FISH

Telomeric DNA was quantified by the incorporation of a nucleic acid telomeric probe (CCCTAA)3 conjugated to Cy5 (TelCy5) by combining flow cytometry with fluorescence in situ hybridization (flow-FISH). PBMCs were stained for 15 min with anti-KLRG1 FITC (REA261, Miltenyi), anti-CCR7 PE (G043H7), anti-CD28 BV786 (CD28.2), anti-CD8 BV605 (SK1), anti-CD27 BV510 (O323), CD45RA BV421 (HI100) from Biolegend and Live/Dead Fixable Near-IR (775) stain (ThermoFisher), after which samples were fixed and permeabilized (Fix & Perm Cell Permeabilisation Kit, Caltag Laboratories). After washing in hybridization buffer (70% deionized formamide, 28·5 m_M_ Tris–HCl pH 7.2, 1·4% BSA and 0·2_M_ NaCl), cells were incubated with 0.75 μg/mL of the PNA telomeric (C3TA2)3 probe conjugated to Cy5 (Panagene). Samples were heated for 10 min at 82°C, rapidly cooled on ice, and hybridized for 1 h at room temperature in the dark. Samples were washed and analysed immediately by flow cytometry.

### Functional Assays

Degranulation was assessed by CD107a externalisation using anti-CD107a APC (H4A3, Biolegend). PBMCs were stimulated with 0.5 µg/µL of anti-CD3 (OKT3) and incubated with anti-CD107a in complete RPMI-1640 media for 45 min at 37°C. 100 µM of monensin (Biolegend) was added and the cells were incubated for a further 4 h and 15 min at 37°C. The cells were washed in PBS and surface and intracellular staining was performed as described above.

Where indicated 10ng/mL TGFβ1 (R&D Systems) was added to complete media and cultured with PBMCs overnight or 5 days as indicated.

In vitro cytotoxicity assays were performed by co-culturing bead-isolated CD8^+^ T cells (Miltenyi), stimulated with 1⍰g/mL anti-CD3 (OKT3, BioLegend), with K562 target cells at the designated effector-to-target (E:T) ratios in complete RPMI-1640 medium for 4⍰h. Prior to co-culture, target cells were labelled with 50⍰µM calcein (Invitrogen) following the manufacturer’s protocol. For each assay, 1 × 10^4^ target cells were incubated with effector cells at E:T ratios ranging from 1.25:1 to 10:1. Additional controls included target cells alone, effector cells alone, and target cells treated with 0.1% Tween-20. Percent specific lysis was calculated using the following formula: % specific lysis = [(experimental lysis – spontaneous lysis)/(maximum lysis – spontaneous lysis)] × 100.

### Quantification of Cytokines

Total IL1-β and TGF-β1 in serum samples were determined either by flow cytometry using the LegendPlex kit (Biolegend) or ELISA (Abbexa) according to the manufacturer’s instructions. To assess the amount of total TGF-β1, acid activation was performed to isolate free TGFβ1 from its latent state as described previously (31).

### Cellular Senescence RT2 Profiler PCR Arrays

Unstimulated CD8^+^ EMRA T cells from T2D and healthy age-matched control participants were isolated using MACS sorting for CD8^+^ T cells and then FACS sorted using anti-CD27/anti-CD45RA for EMRA isolation. RNA was extracted using the RNAeasy Micro Kit (Qiagen) and pre-amplified using the RT2 PreAMP cDNA Synthesis Kit (Qiagen) and RT2 PreAMP Pathway Primer Mix (Qiagen). The resulting gene-specific cDNA was then analysed using the Cellular Senescence RT2 Profiler PCR Arrays according to the manufacture’s instructions (Qiagen).

### qPCR of senescence markers

RNA from MACS (Miltenyi Biotec) purified CD8+ T cells were isolated using the RNeasy kit (Qiagen) according to the manufacturer’s instructions. Transcripts were quantified using the High-Capacity cDNA Reverse Transcription Kit (Applied Biosystems) and the SsoAdvanced Universal SYBR Green Supermix (Bio-Rd Laboratories) according to the manufacturer’s instructions. The following primers were purchased from IDT.

TGFBR1 F: GCT GTA TTG CAG ACT TAG GAC TG; TGFBR1 R: TTT TTG TTC CCA CTC TGT GGT T; TGFBR2 F: AAG ATG ACC GCT CTG ACA TCA; TGFBR2 R: CTT ATA GAC CTC AGC AAA GCG AC; TGFBR3 F: TGG GGT CTC CAG ACT GTT TTT; TGFBR3 R: CTG CTC CAT ACT CTT TTC GGG; p53 F: GCC AAG TCT GTG ACT TGC ACG; p53 R: TGT GGA ATC AAC CCA CAG CTG, p21 F: GGC AGA CCA GCA TGA CAG ATT TC; p21 R: CGG ATT AGG GCT TCC TCT TGG, p16 F: CAA GAT CAC GCA AAA ACC TCT G, p16 R: CGA CCC TAT ACA CGT TGA ACT G, DAB2 F: GTA GAA ACA AGT GCA ACC AAT GG, DAB2 R: GCC TTT GAA CCT TGC TAA GAG A, rab11a F: CAA CAA GAA GCA TCC AGG TTG A, rab11a R: GCA CCT ACA GCT CCA CGA TAA T rab27a F: ACA ACA GTG GGC ATT TTC A, rab27a R: AAG CTA CGA AAC CTC TCC TGC, BActin F: CAC CAT TGG CAA TGA GCG GTT C; BActin R: AGG TCT TTG CGG ATG TCC ACG T.

### TCR repertoire analysis

Unstimulated CD8^+^ EMRA T cells from 3 T2D and 3 healthy age-matched participants were isolated using CD8 MACS positive selection followed by FACS sorting using anti-CD27/anti-CD45RA for EMRA isolation. The resulting EMRA T cells were used to create a TCR library according to previously published protocols (32). The data was analysed using Decombinator V3 and V5, a software package which allows for fast, efficient gene assignment in T cell receptor sequences using a finite state machine (33). The resulting output was analysed further using R. Diversity indexes, such as Gini-coefficient and Shannon’s entropy were analysed after bootstrapping.

### ^89^Zirconium radiolabelling and PET imaging

[^89^Zr]Zr(oxalate) (Perkin Elmer) was chelated with oxine forming [^89^Zr]Zr(oxinate)_4_ as previously described(34). CD8+ T cells were isolated by positive selection using the MACS system (MiltenyiBiotec) according to the manufacturer’s instructions and were mixed with [^89^Zr]Zr(oxinate)_4_ with a volume ratio not lower than 30:1 (cells: [^89^Zr]Zr(oxinate)_4_). Radioactivity was counted using a Wallac Wizard gamma counter to determine cell-associated radioactivity and the percentage of injected activity (%IA) present in cells was then calculated.

^89^Zr-labelled CD8+ T cells were injected intravenously into female Nod *scid* gamma (NSG, NOD.Cg-*Prkdc* ^scid^ *IL2rg*^*tm1Wjl*^/SzJ) mice aged 6 weeks, average body weight of 20.7⍰±⍰1.5⍰g (Charles River) at a concentration of 3×10^6^cell/animal under anaesthesia (1.5-2.5% isoflurane in oxygen). Female NSG mice were used in this study as they were found to better support engraftment of human immune cells (35). Animal experiments were approved by the UK Home Office under The Animals (Scientific Procedures) Act (1986), with local approval from King’s College London Animal Welfare and Ethics Review Body. All procedures complied with relevant guidelines and regulations and have been reported in accordance with the ARRIVE guidelines.

Three hours post-injection, mice were re-anaesthetised and placed in a preclinical nanoPET/CT scanner (Mediso) where anaesthesia was maintained and the bed was heated to maintain a normal body temperature. Two hours of PET acquisition (1:5 coincidence mode; 5 ns coincidence time window) were followed by CT. PET-CT was repeated at t = 24 and 72 h. PET/CT images were reconstructed using a Monte Carlo-based full-3D iterative algorithm (Tera-Tomo, 400-600 keV energy window, 1-3 coincidence mode, 4 iterations and subsets) at a voxel size of (0.4 × 0.4 × 0.4) mm^3^ and corrected for attenuation, scatter, and decay. Images were co-registered and analysed using VivoQuant v.3.0 (InVicro LLC) capturing maximum intensity projection (MIP) and transverse plane images. If required, background, not associated with anatomy, was manually removed. Spherical VOIs were placed on organs based on the CT image when the spleen tissue was not visible. %IA was calculated to assess overall injected activity in each animal and %IA/mL to determine injected activity distribution and concentration in selected organs.

Mice from imaging studies were used for biodistribution studies at 72 h post injection. After culling, organs were dissected, weighed, and γ-counted together with standards prepared from a sample of injected material. The percentage of injected activity per gram injected dose per gram (%IA/g) of tissue was calculated.

### Plaque digestion

Atrial plaques were processed within 2 h of surgery, and digested with 300U/ml Collagenase Type IV (Sigma) and 300U DNase I (Sigma) for 30 min at 37°C. The released cells were strained first through a 100µm then a 40µm cell strainer. CD3+ T cells were isolated by MACS positive selection (MiltenyiBiotec) according to the manufacturer’s instructions. The resulting T cells were then phenotyped by flow cytometry.

### Statistical analysis

GraphPad Prism was used to perform statistical analysis. Statistical significance was evaluated using a Mann-Whitney U test or ANOVA followed by Tukey multi-comparison test. Diversity within T cell repertoire data was assessed by calculating the Shannon Diversity Index and dispersion with the Gini index. Both indices were computed using R and compared across conditions using a Mann-Whitney U test. Graphs show SD. Differences were considered significant when P was <0.05. Stars signify the following: *p<0.05, **p<0.01, ****p<0.0001.

### Code Availability

No new code was used to generate the data. TCR analysis was performed using the Decombinator package (36), while the Circos package (37) was used to generate the Circos plots.

## Results

### Heterogeneity of CD8+ TEMRA cells in healthy ageing and T2D-associated unhealthy immune ageing

CD8^+^ T cell subsets are classified based on their function, location and expression of surface markers, with T cell differentiation often being defined based on CD45RA and CCR7 expression (19). Using these markers CD8^+^ T cells can be divided into naïve (T_N_ ; CD45RA^+^CCR7^+^), central memory (T_CM_ ; CD45RA^-^CCR7^+^), effector memory (T_EM_; CD45RA^-^CCR7^-^) and effector memory CD45RA re-expressing cells (T_EMRA_ ; CD45RA^+^CCR7^-^). We initially employed these markers to quantify terminally differentiated T cells in older adults with T2D (>55 years), with T2D serving as the primary model of T2D-associated unhealthy immune ageing (Supplementary Figure 1A). The study cohorts were matched for age, sex, BMI, CMV serostatus and ethnicity (Table 1). Consistent with this, T_EMRA_ frequency was not influenced by participant gender (Supplementary Figure 1B) or ethnicity (Supplementary Figure 1C). Moreover, stratification by HbA1c revealed no differences in T_EMRA_ frequencies, indicating that the observed phenotype in T2D is not explained by glycaemic control alone supporting its use as a model of premature immune ageing (Supplementary Figure 1D). In line with previous reports, we found a higher percentage of CD45RA/CCR7 defined T_EMRA_ cells in participants with T2D (Figure 1A). However, these markers fail to capture the heterogeneity within the T_EMRA_ subset. Therefore, we further defined differentiation using the co-stimulatory receptors CD27 and CD28, which resulted in three populations increasing in differentiation from CD27^+^CD28^+^ (E1) CD27^+^CD28^-^ (E2) through to CD27^-^CD28^-^ (E3) cells. Incorporating these markers revealed that participants with T2D exhibited a significantly higher proportion of highly differentiated E3 cells (Figure 1B).

**Figure 1.**
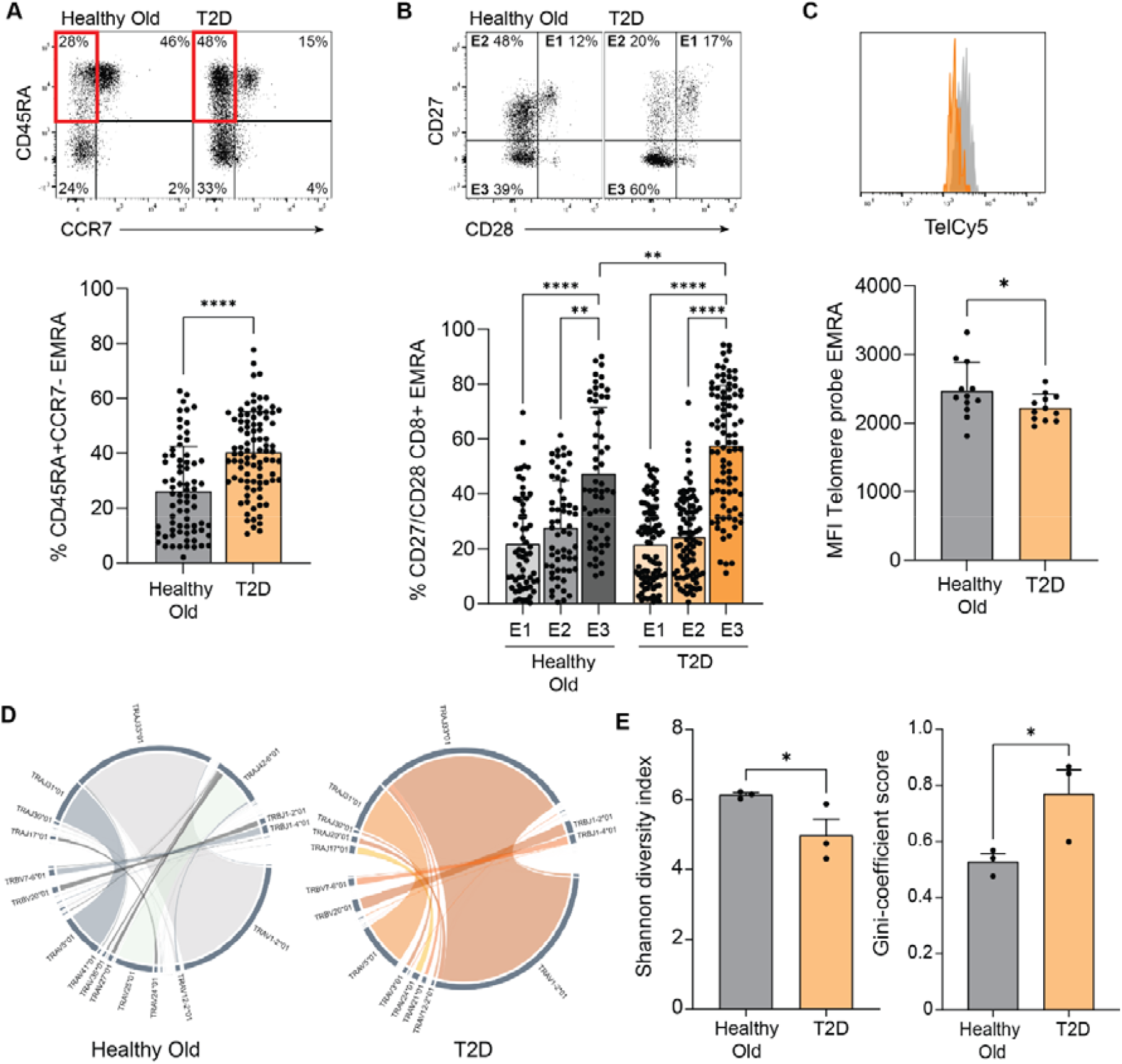
CD8^+^ T_EMRA_ cells in T2D-associated unhealthy immune ageing are highly differentiated. (A) Flow cytometry plot and graph showing CD45RA/CCR7 defined CD8^+^ T cell populations. The red box denotes the CD45RA^+^CCR7^-^ T_EMRA_ cells. (B) Flow cytometry plot and graph showing CD27/CD28 expression in the CD45RA^+^CCR7^-^ T_EMRA_ population. (C) Histogram and graph showing telomere length in CD45RA/CCR7 defined T_EMRA_ cells. (D) Chord diagrams displaying the frequencies and clonalities of the TCR from T_EMRA_ cells. (E) Graphs showing Shannon diversity index and Gini-coefficient values for CD8^+^ T_EMRA_ populations. All data compares individuals with and without T2D, was analysed using a Mann-Whitney U test and is expressed as mean ± SD. *p<0.05, **p<0.01 and ****p<0.0001.

To confirm that the increase in highly differentiated T_EMRA_ cells was not driven by high glucose levels, we repeated our analysis on a separate cohort of individuals with dilated cardiomyopathy (DCM), comparing those with and without T2D. We observed that individuals with cardiomyopathy also had a greater amount of CD45RA/CCR7 defined T_EMRA_ cells, with no significant difference between the T2D and non-T2D groups (Supplementary Figure 2A). Furthermore, these T_EMRA_ cells from unhealthily aged individuals contained a higher proportion of highly differentiated E3 cells (Supplementary Figure 2B). Together, these findings suggest that the observed CD8^+^ T_EMRA_ populations should not be attributed solely to immune ageing but are associated with inflammatory and metabolically dysregulated disease contexts that may overlap with features of unhealthy immune ageing.

A functional assessment of differentiation further confirmed the highly differentiated nature of T_EMRA_ cells in participants with T2D. Telomere length analysis revealed significant telomere shortening in these cells, indicating increased replicative history and potential senescence (Figure 1C). Additionally, TCR sequencing in individuals with T2D compared to healthy older adults demonstrated a shift towards a more oligoclonal repertoire in the T2D group (Figure 1D, Supplementary Figure 2C), characterised by reduced diversity and the disproportionate expansion of a few dominant T cell clones (Figure 1E). These findings are consistent with enhanced terminal differentiation and expansion of T_EMRA_ cells in this context, however the small sample size means they should be interpreted with caution and require validation in a larger cohort.

Interestingly, despite their highly differentiated phenotype, T_EMRA_ cells from individuals with T2D showed reduced expression of markers traditionally associated with highly differentiated T cells, including KLRG1 and the NK receptors NKG2D and NKG2A, suggesting a divergence from the expected characteristics of highly differentiated CD8^+^ T cells. KLRG1 an inhibitory receptor commonly associated with replicative senescence and reduced proliferative capacity in T cells was found to be significantly lower in T_EMRA_ cells from individuals with T2D (Figure 2A). Similarly, both the activatory NKG2D and inhibitory NKG2A receptors were also reduced (Figure 2B). Analysis of the internal-to-surface KLRG1 ratio revealed a higher proportion of intracellular KLRG1 within T_EMRA_ cells from individuals with T2D (Figure 2C), consistent with altered receptor trafficking. This phenomenon was consistent across individuals with cardiomyopathy, further supporting the finding (Supplementary Figure 2D). To investigate the mechanisms underlying this trafficking defect, qPCR analysis was performed on key regulators of receptor internalisation and recycling. Expression of *dab2*, an adaptor involved in clathrin-mediated endocytosis (38), was unchanged in CD8^+^ T cells from individuals with T2D suggesting that receptor internalisation is not impaired (Figure 2D). In contrast, expression of *rab11a*, a small GTPase that regulates endosomal recycling and the return of internalised receptors back to the plasma membrane (39), was significantly reduced (Figure 2D). This reduction is consistent with impaired receptor recycling, leading to intracellular accumulation of KLRG1 due to inefficient trafficking, although increased degradation cannot be excluded the data favour a defect in receptor recycling.

**Figure 2.**
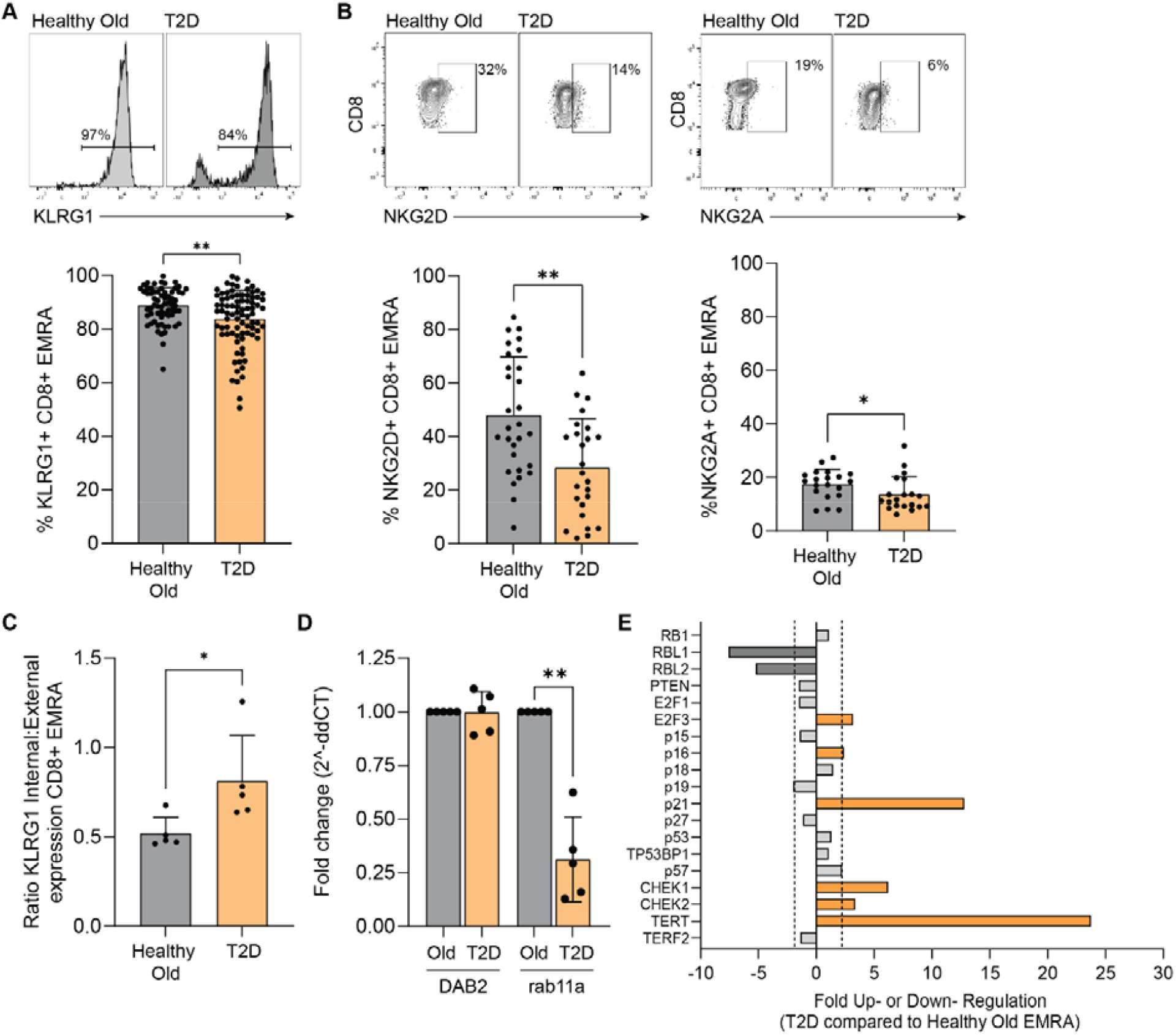
CD8^+^ T_EMRA_ cells in T2D-associated unhealthy immune ageing show altered localisation of phenotypic markers associated with senescence. (A) Histogram and graph showing expression of KLRG1 in CD45RA/CCR7 defined CD8^+^ T_EMRA_ cells. (B) Flow cytometry plots and graphs showing expression of NKG2D and NKG2A in CD45RA/CCR7 defined CD8^+^ T_EMRA_ cells. (C) Ratio of internal: external expression of KLRG1 in T_EMRA_ cells. (D) Relative mRNA expression of *dab2* and *rab11a* in CD8^+^ T_EMRA_ cells from people with or without T2D, measured by qPCR. Expression was normalised to βActin and calculated using the ΔΔCt method. (E) Graph showing senescence-associated genes in T_EMRA_ cells that were up or down-regulated >⍰2-fold using an RT^2^ profiler PCR array, n=3 for each group. Genes with a fold change ≥ 2 were considered significant. All data compares individuals with and without T2D, was analysed using a Mann-Whitney U test and is expressed as mean ± SD. *p<0.05 and **p<0.01.

Interestingly, KLRG1 is a marker commonly used to define tissue-resident memory T cells (T_RM_), and T_EMRA_ cells from both healthy older participants and individuals with T2D expressed lower levels of CD103 and CD69 compared to the overall CD8^+^ T cell population. However, CD8^+^ T_EMRA_ cells from individuals with T2D showed higher CD103 and CD69 expression than those from healthy individuals, indicating a more tissue-resident-like phenotype (Supplementary Figure 2E). These findings suggest that in the T2D-associated disease context, T_EMRA_ cells undergo an altered regulatory process that may affect their function and responsiveness to external stimuli.

The altered phenotype of T_EMRA_ cells suggests that differentiation in T2D-associated unhealthy immune ageing may be shaped by multiple signals rather than a single pathway. We therefore used a Cellular Senescence RT2 Profiler Array to examine whether inflammatory cues contributed to this phenotype within the broader context of metabolic dysfunction. The senescence-pathway array showed increased expression of p21-associated transcripts without a corresponding increase in p53 transcript expression in CD8^+^ T_EMRA_ cells from people with T2D (Figure 2E), a pattern consistent with p21-associated signalling but insufficient to establish a p53-independent mechanism. Although p53 transcript levels did not change, we cannot exclude a role for p53 in regulating p21, as p53 activity is primarily controlled post-translationally (40) meaning that direct modulation of p53 would be required to definitively assess its contribution.

Additionally, one of the most striking observations from the senescence array was the enhanced expression of *TERT*, the catalytic subunit of telomerase. *TERT* was significantly upregulated in T_EMRA_ cells from individuals with T2D (Figure 2E). This increase in *TERT* suggests that T_EMRA_ cells may be relying on non-telomere-dependent mechanisms to sustain cell viability and function under stress, potentially compensating for the observed telomeric attrition.

### TGFβ1 recapitulates features of the TEMRA phenotype associated with T2D-related unhealthy immune ageing

The data presented so far highlight the presence of a CD8^+^ T_EMRA_ population with reduced KLRG1 expression in T2D-associated unhealthy immune ageing, where these cells display features consistent with premature senescence rather than a conventional replicative-senescence phenotype. We have published previously that sera from individuals with T2D contained elevated levels of inflammatory mediators capable of inducing senescence(29). Within this inflammatory secretome we were particularly interested in TGFβ1, given its ability to induce senescence in a p53-independent but p21-dependent manner (41) and because circulating TGFβ1 has been linked to T2D and metabolic dysfunction (42). We therefore investigated whether TGFβ1 may contribute to the emergence of these KLRG1-low T_EMRA_ cells in the T2D-associated disease context.

TGFβ1 levels were significantly higher in individuals with T2D (Figure 3A, Supplementary Figure 3A), and its presence correlated with the accumulation of T_EMRA_ cells (Supplementary Figure 3B). Furthermore, T_EMRA_ cells from individuals with T2D exhibited higher expression of TGFβ receptor 3 (TGFβR3) (Supplementary Figure 3C). In addition to the increased expression of TGFβ1 in T2D-associated T_EMRA_ cells, the RT2 Profiler Array also revealed an upregulation of TGFβ1i1 (transforming growth factor beta 1 induced transcript 1), which enhances TGFβ1 signalling through the inhibition of Smad7 (43) (Supplementary Figure 3D). While we did not directly assess the cellular source of TGFβ1 our data support a model in which systemic inflammation, rather than cell-intrinsic production of TGFβ1 exerts its effect on T_EMRA_ cells. Interestingly, a significant proportion of participants were taking statins, which are known to reduce TGFβ1 levels (44, 45). When we stratified the data from individuals without T2D, where there was a more even split between those taking and not taking statins, we found that those on statins had a lower proportion of T_EMRA_ cells (Supplementary Figure 3E). However, while statin use was associated with a reduction in serum TGFβ1 levels in healthy individuals, this effect was not observed in individuals with T2D, (Supplementary Figure 3F), suggesting that the inflammatory environment in T2D may override the TGFβ1-lowering effects of statins.

**Figure 3.**
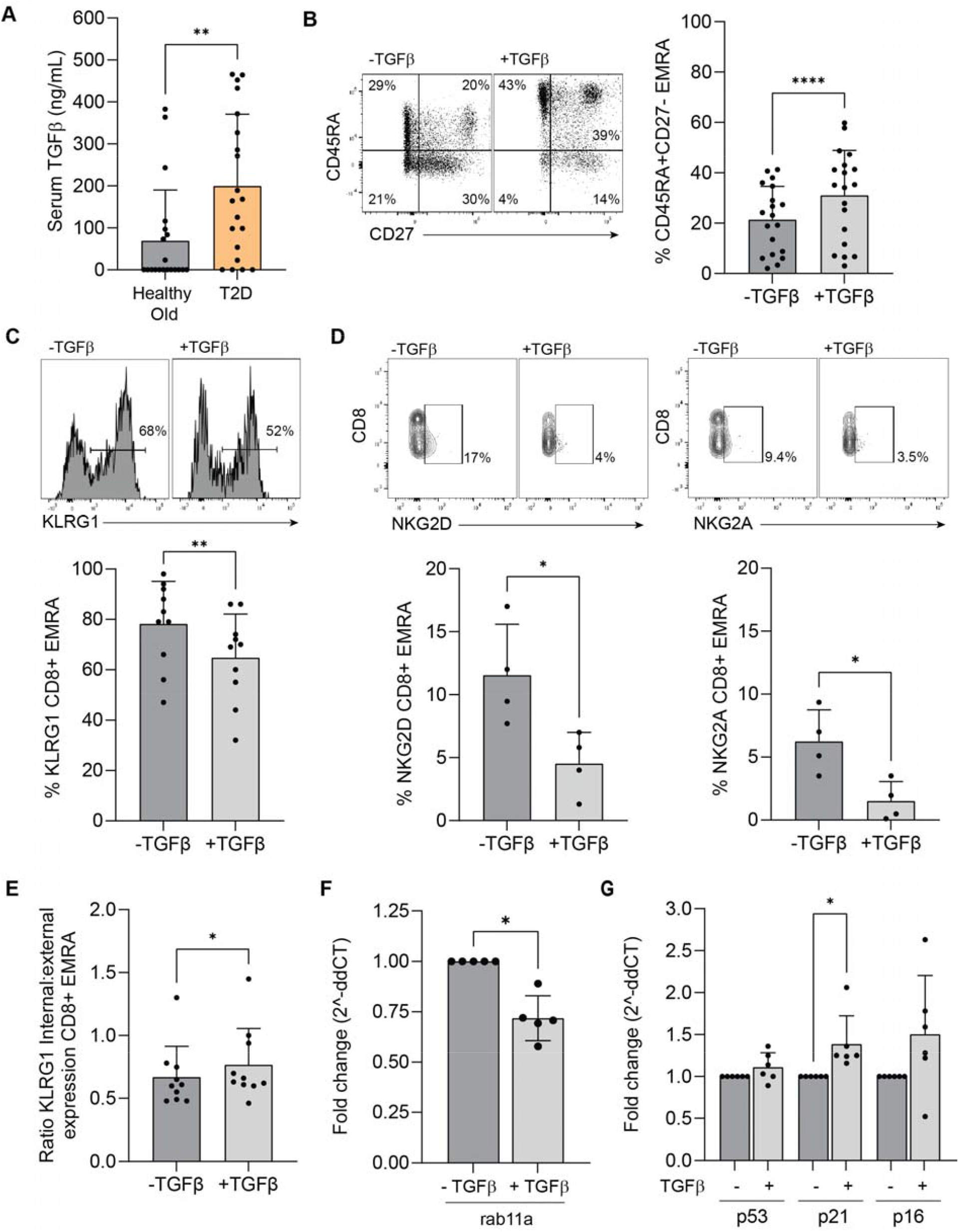
TGFβ1 recapitulates features of the CD8^+^ T_EMRA_ phenotype associated with T2D-related unhealthy immune ageing. (A) Serum TGFβ1 levels measured by ELISA in individuals with and without T2D. (B) Flow cytometry plot and graph showing CD45RA/CCR7 defined CD8^+^ T cell populations with and without 10ng/mL TGFβ1. (C) Histogram and graph showing expression of KLRG1 in T_EMRA_ cells with and without TGFβ1. (D) Flow cytometry plots and graphs showing expression of NKG2D and NKG2A in CD45RA/CCR7 defined CD8^+^ T_EMRA_ cells with and without TGFβ1. (E) Ratio of internal: external expression of KLRG1 in T cells with and without TGFβ1. (F) Relative mRNA expression of *rab11a* in CD8^+^ T cells incubated with and without TGFβ1 for 5 days. Expression was normalised to βActin and calculated using the ΔΔCt method. (G) Relative gene expression of p53, p21 and p16 using RT-PCR in T_EMRA_ cells with and without TGFβ1. All data was analysed using a Mann-Whitney U test and is expressed as mean ± SD. *p<0.05, **p<0.01 and ****p<0.0001.

To determine whether TGFβ1 could induce a premature T phenotype, we incubated CD8^+^ T_EMRA_ cells with 10 ng/ml TGFβ1 and assessed their differentiation. TGFβ1 treatment led to a significant increase in T_EMRA_ cells (Figure 3B), accompanied by phenotypic changes observed in the T2D cohort, including reduced KLRG1 expression (Figure 3C) as well as lower levels of both NKG2D and NKG2A (Figure 3D). Incubation with TGFβ1 also increased the ratio of intracellular to extracellular KLRG1 levels (Figure 3E) and was accompanied by reduced expression of *rab11* (Figure 3F). Additionally, TGFβ1 increased p21 transcript abundance without increasing p53 transcript abundance (Figure 3G), consistent with a p21-associated response but insufficient to establish p53-independent signalling. These findings identify TGFβ1 as a candidate contributor to several components of the premature senescence-like T_EMRA_ phenotype observed in T2D, potentially linking inflammatory and metabolic cues to altered CD8+ T cell differentiation.

### T_EMRA_ cells in in T2D-associated unhealthy immune ageing are associated with functional impairment

TGFβ can influence adhesion by modulating the expression of chemokine receptors, cell adhesion molecules and by influencing extracellular matrix (ECM) synthesis and remodelling (46, 47); we chose to examine here tissue distribution and ECM-associated pathways. To determine whether elevated TGFβ1 levels contribute to altered trafficking and tissue residency of T_EMRA_ cells in T2D-associated unhealthy immune ageing, we investigated its impact on distribution patterns and integrin expression.

We isolated CD8^+^ T cells from both healthy older adults and individuals with T2D and radiolabelled them with zirconium-89 ([^89^Zr]Zr-(oxinate)_4_). These ^89^Zr-labelled CD8^+^ T cells were then injected into NOD scid gamma (NSG) mice at an average activity of 24 ± 18 kBq, and their movement was tracked using PET imaging. The same concentration of CD8^+^ T cells was administered for both source groups. Previous work using this radiolabelling approach has shown that ^89^Zr-labelled CD8^+^ T cells remain viable and intact after labelling, with no increase in cell death supporting the use of this method for tracking labelled T cells in vivo (48). PET scans revealed a significantly greater accumulation of ^89^Zr-labelled CD8^+^ T cells from individuals with T2D (Figure 4A), whereas CD8^+^ T cells from individuals without T2D exhibited a more diffuse distribution, resulting in signal levels below the sensitivity threshold of the PET imaging system. While these experiments were conducted in lymphopenic NSG mice to prevent alloreactivity, the host environment was consistent across all source cell types. Accordingly, differences between donor groups are consistent with source-cell-dependent intrinsic or compositional properties of the CD8^+^ T cell preparations under equivalent recipient conditions, rather than differences in the recipient host.

**Figure 4.**
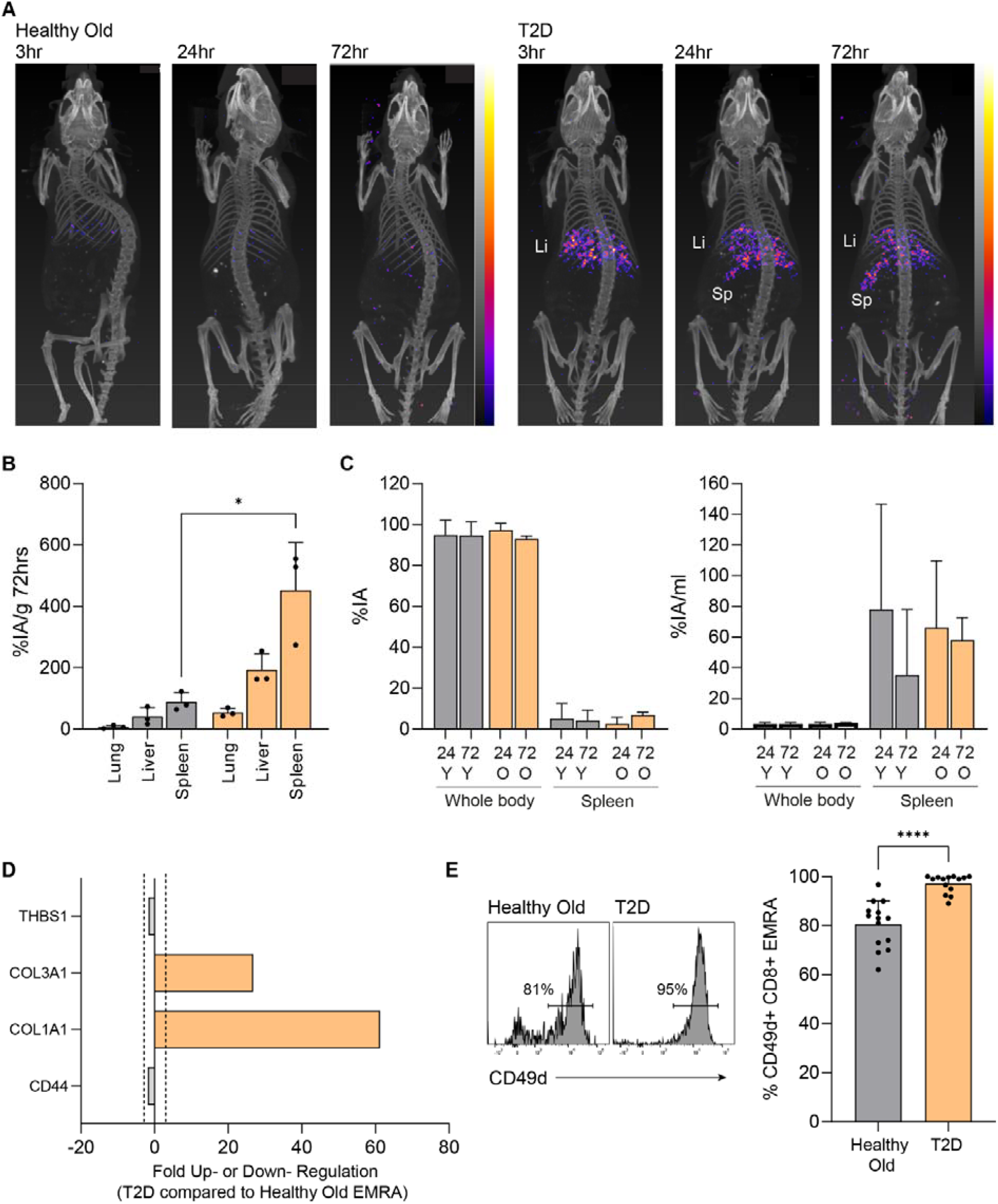
T2D-derived CD8^+^ T cells show altered tissue distribution and T_EMRA_ cells show an adhesion-associated phenotype. (A) Representative images of NSG mice injected intravenously with ^89^Zr-labelled 3 × 10^6^ CD8^+^ T cells from individuals with and without T2D. Mice were subjected to whole-body preclinical PET at 3 h, 24 h and 72 h. (B) *Ex vivo* biodistribution of ^89^Zr-labelled CD8+ T_EMRA_ cells isolated from individuals with and without T2D 72 h after CD8^+^ T cell administration. (C) Graphs showing the image-based analysis of percentage injected activity (%IA) and injected activity/mL, calculated for the whole-body and spleen tissue VOIs of 2 mice per group at 24 and 72 h after the administration of radioactive CD8^+^ T cells. (D) Graph showing adhesion-related genes in T_EMRA_ cells that were up-regulated >⌇2-fold using an RT^2^ profiler PCR array. Genes with a fold change ≥ 2 were considered significant. Data compares individuals with and without T2D, n = 3 for each group. (E) Histogram and graph showing expression of CD49d in T_EMRA_ cells from individuals with and without T2D. Data was analysed using a Mann-Whitney U test and is expressed as mean ± SD. *p<0.05 and ****p<0.0001.

Biodistribution analysis showed that the highest concentration of ^89^Zr-labelled CD8^+^ T cells was found in the spleen, with a significantly higher accumulation of CD8^+^ T cells from individuals with T2D compared to those from healthy older individuals (Figure 4B). To further examine the dynamics of this accumulation, we performed serial PET imaging and quantified the percentage injected activity (%IA) over time (Figure 4C). This analysis confirmed that while most of the radioactivity was retained within the whole-body volume of interest (VOI), the temporal accumulation profiles in the spleen were comparable between groups, with no clear evidence of accelerated initial accumulation of T2D-derived CD8^+^ T cells. Instead, the higher splenic accumulation is consistent with altered retention or preferential homing of T2D-derived CD8^+^ T cells influenced by the inflammatory environment in the context of unhealthy ageing. Although this model allows comparison of relative trafficking differences between CD8^+^ T cells from healthy and T2D donors under equivalent recipient conditions, it does not fully recapitulate a non-lymphopenic T cell compartment or the inflammatory and metabolic environment of T2D. Therefore, these findings should be interpreted as preliminary evidence of altered trafficking potential.

This shift in trafficking dynamics may be linked to collagen remodelling, a process regulated by TGFβ (49). We show that CD8^+^ T cells from individuals with T2D exhibited increased expression of collagen genes, COL1A1 and COL3A1, (Figure 4D) and the integrin CD49d (Figure 4E), consistent with an enhanced adhesion-associated phenotype. Additionally, incubation with TGFβ1 increased CD49d expression in CD8^+^ T_EMRA_ cells (Supplementary Figure 4A).

To examine whether the altered phenotype was also evident in human tissue, we quantified T_EMRA_ cells in paired peripheral blood and carotid atherosclerotic plaques (Figure 5A), where they exhibited a premature senescent phenotype characterised by reduced KLRG1 expression (Figure 5B). Indicating differential representation of T_EMRA_ cells in plaques.

**Figure 5.**
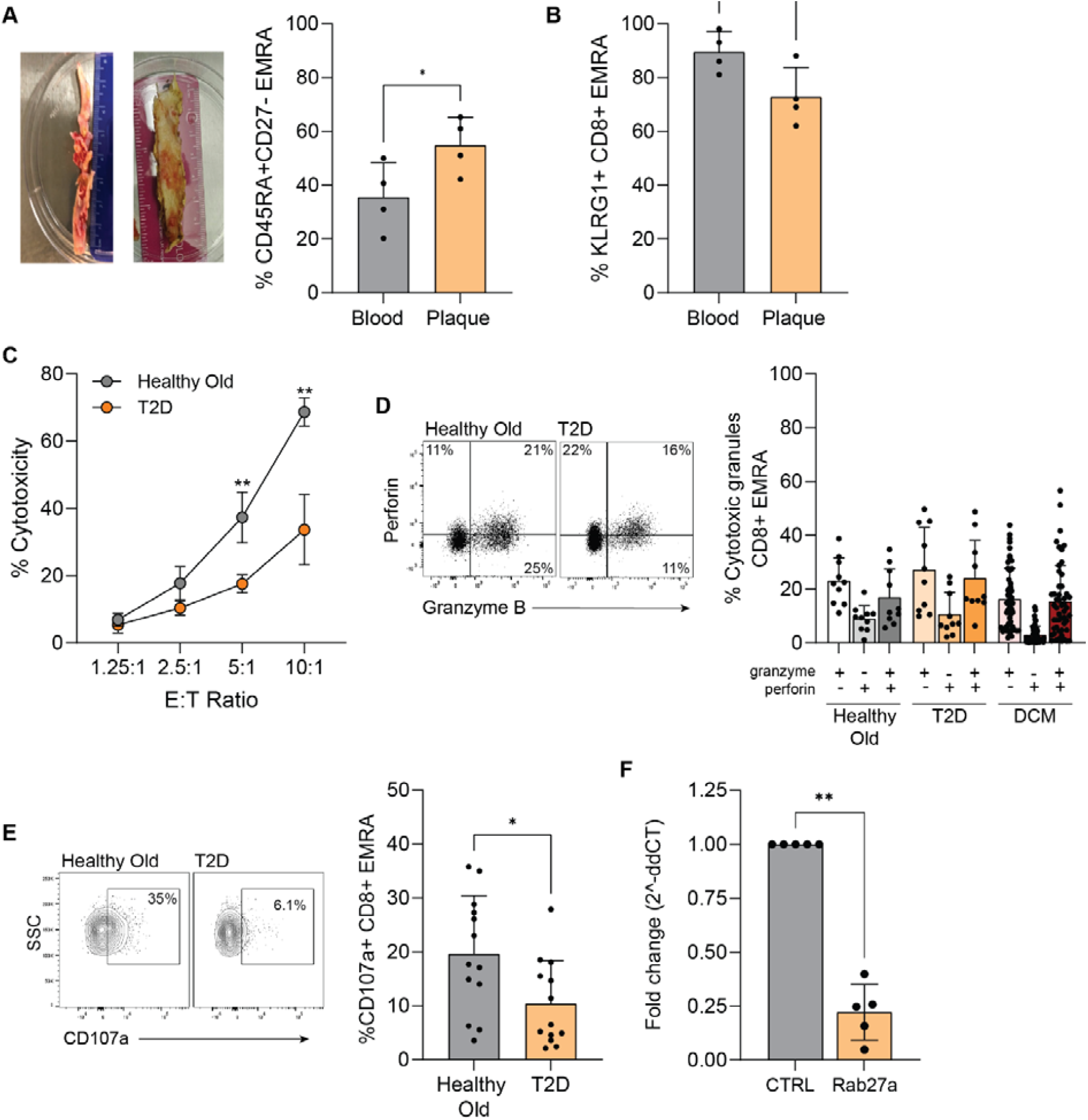
CD8^+^ T_EMRA_ cells in T2D-associated unhealthy immune ageing show altered plaque representation and reduced stimulated effector responses. (A) Representative atherosclerotic plaques and paired data showing the percentage of total CD8^+^ T cells with a CD45RA^+^CCR7^-^ T_EMRA_ phenotype in peripheral blood and plaque, n=4. (B) Percentage of KLRG1^+^ cells within the T_EMRA_ subset in paired peripheral blood and plaque samples. (C) Specific lysis/cytotoxicity of calcein-labelled K562 target cells by soluble anti-CD3-stimulated, bead-isolated total CD8^+^ T_EMRA_ cells from healthy older participants or individuals with T2D, n=3 per group. (D) Flow cytometry plot and graph showing granzyme B and perforin expression in the CD45RA^+^CD27^-^ T population in healthy older participants, individuals with T2D and individuals with dilated cardiomyopathy. (E) Flow cytometry plots and graphs showing CD107a externalisation within the phenotypically defined CD8^+^ T_EMRA_ population following anti-CD3 stimulation in healthy older participants and individuals with T2D. This assay measures TCR/CD3-stimulated degranulation. (F) Relative mRNA expression of rab27a in bulk CD8^+^ T cells from individuals with or without T2D. Expression was normalised to βActin and calculated using the ΔΔCt method. Data were analysed using a Mann-Whitney U test and are expressed as mean ± SD. *p<0.05, **p<0.01.

Despite showing similar levels of the cytotoxic granules perforin and granzyme B (Figure 5C), T_EMRA_ cells from individuals with unhealthy ageing exhibited significantly reduced CD107a expression (Figure 5D), indicating a defect in degranulation rather than granule production. This impaired degranulation was found to be TGFβ1-mediated, as incubation of T_EMRA_ cells with TGFβ1 from healthy individuals induced a similar decrease in CD107a expression (Supplementary Figure 2G). Furthermore, TGFβ induced signalling has been shown to suppress CD69 expression on numerous immune cells (50). Consistent with this, we observed a downregulation of CD69 expression on T_EMRA_ cells from individuals with unhealthy ageing (Figure 5E). This reduction in CD69 may contribute to an altered activation state of T_EMRA_ cells, potentially promoting their retention in secondary lymphoid organs. As a result, this dysregulated activation may impair their ability to respond effectively to external stimuli, ultimately decreasing their activation and overall immune function.

Functional assays were used to determine whether the altered phenotype of T_EMRA_ cells in T2D-associated unhealthy immune ageing was accompanied by impaired effector responses. Anti-CD3-stimulated bulk CD8^+^ T cells from individuals with T2D showed reduced specific lysis of K562 targets compared with cells from healthy older participants (Figure 5C). As K562 cells are highly susceptible to NK-like cytotoxic mechanisms, this finding is consistent with reduced cytotoxic competence in the context of the lower NK-receptor expression observed in T2D-associated T_EMRA_ cells, although the use of anti-CD3 stimulation means that this assay does not directly measure NK-receptor-mediated or TCR-independent killing. Despite broadly similar expression of the cytotoxic granules granzyme B and perforin within the T_EMRA_ population (Figure 5D), phenotypically defined T_EMRA_ cells from individuals with T2D exhibited reduced CD107a externalisation following anti-CD3 stimulation (Figure 5E), consistent with impaired degranulation rather than reduced granule abundance. This phenotype was accompanied by reduced *rab27a* transcript abundance in bulk CD8^+^ T cells (Figure 5F), consistent with altered regulation of the vesicular trafficking machinery required for granule exocytosis. TGFβ1 treatment of T_EMRA_ cells from healthy individuals similarly reduced CD107a expression, suggesting that TGFβ1 may contribute to the impaired degranulation phenotype observed in T2D.

Consistent with this phenotype, individuals with T2D reported poorer overall health and greater pain (Supplementary Figure 5A,B), and T_EMRA_ frequency positively correlated with increased IL-1β production (Supplementary Figure 5C). These questionnaire data provide clinical context for the T2D cohort, without implying a direct causal link between patient-reported health and CD8^+^ T_EMRA_ frequency or function. Together, the data are consistent with a dysregulated T_EMRA_ state in T2D-associated unhealthy immune ageing, characterised by altered tissue representation and reduced TCR-stimulated effector responses. Whether this phenotype contributes directly to tissue damage or chronic inflammation remains to be determined.

## Discussion

Ageing is accompanied by significant changes in immune function, but the trajectory of these changes differs between healthy and unhealthy ageing. In healthy ageing, the immune system undergoes gradual adaptations while maintaining sufficient responsiveness to infections and immune challenges. In contrast, unhealthy ageing is characterized by excessive immune dysregulation, including chronic inflammation and metabolic stress (51). Using T2D as the primary model of disease-associated unhealthy immune ageing, we identify a population of CD8^+^ T_EMRA_ cells that downregulate conventional phenotypic markers of T cell senescence and instead show features consistent with premature senescence, with TGFβ recapitulating several components of this phenotype in vitro. These T2D-associated T_EMRA_ cells show a more tissue-resident-like phenotype and reduced degranulation following anti-CD3 stimulation, consistent with altered effector responsiveness.

An important mechanism for preserving immune protection in older individuals is the enhanced expression of NK receptors on CD8^+^ T_EMRA_ cells, which help maintain their cytotoxic capacity and compensate for the decline in T cell repertoire diversity (52). However, we identify a subset of CD8^+^ TEMRA cells in T2D-associated unhealthy immune ageing that show reduced NK receptor expression. While our functional data addresses whether this phenotypic change impairs innate-like cytotoxicity, a direct functional link remains to be fully established. Mechanistically, this phenotype appears to be associated with reduced Rab expression. Rab GTPases are central regulators of endosomal trafficking, receptor recycling to the plasma membrane and lytic granule exocytosis (53). Disruption of these pathways leads to intracellular retention of key surface receptors and impaired cytotoxic function. Our findings contrast with a study that reported an age-related increase in granzyme K (GZMK)-expressing CD8^+^ effectors (54). However, this study was conducted in healthy, lean older adults, suggesting that its conclusions may not fully reflect the immune alterations seen in unhealthy ageing.

Despite the loss of NK receptors, CD8^+^ TEMRA cells in T2D-associated unhealthy immune ageing were found to be highly differentiated both phenotypically and functionally, exhibiting shorter telomeres and a more oligoclonal repertoire than T_EMRA_ cells from individuals undergoing healthier ageing. Altered systemic CD8^+^ T cell and NK cell proportions have been reported in T2D patients, supporting the concept that immune composition is reshaped in metabolic disease (55). A decline in TCR diversity is typically driven by homeostatic expansion of the peripheral T cell pool following lymphopenia, often in response to IL-7 and IL-15 (56). However, homeostatic expansion can also be triggered by self-antigens, contributing to the development of autoimmunity (57). Indeed, we show here that T_EMRA_ cells from individuals with T2D exhibited a higher frequency of publicly expanded TCR sequences, particularly those utilising TRAV1-2. This TCR bias gives the repertoire a germline-like quality, characterised by low diversity, high generation probability and frequent publicness. Such features are commonly associated with TCRs recognising conserved microbial antigens, suggesting that chronic exposure to microbial or microbiota-derived antigens may shape the TCR repertoire (58). As chronic antigen exposure depends on MHC-I presentation, glucose- and metabolism-driven changes in MHC-I expression may alter antigenic and tonic signals received by CD8^+^ T cells (59, 60). The enrichment of public TCRs also raises the possibility of shared, antigen-driven immune responses that could contribute to the development or perpetuation of inflammation in the T2D-associated disease context.

Interestingly, telomerase reverse transcriptase (TERT) the catalytic subunit of the enzyme telomerase was found to be higher in CD8+ T_EMRA_ cells from individuals with T2D, despite these cells having shorter telomeres. This suggests that TERT may play an extra-telomeric role in this context (61). Telomere-independent activities of TERT have been shown to influence many essential cellular processes, such as gene expression, signalling pathways, mitochondrial function and cell survival (62-64). Additionally, TERT can interact with the NF-κB pathway contributing to inflammation (65), which may further support the inflammatory secretome associated with CD8^+^ T_EMRA_ cells in T2D-associated unhealthy immune ageing(29).

Our study suggests that CD8^+^ T_EMRA_ cells are associated with exposure to TGFβ1, which recapitulates several features of the phenotype in vitro. TGFβ signalling is important both for immunity, where it controls the differentiation of CD8^+^ T cells, as well as regulating senescence (66). The role of TGFβ in senescence has been well documented; being involved in cell proliferation, cell cycle regulation, the production of reactive oxygen species (ROS), DNA damage repair, telomere regulation, unfolded protein response (UPR), and autophagy (67). Importantly, TGFβ is a potent inducer of premature senescence activating the Smad pathway responsible for upregulating p21 (68). It is also secreted as part of the inflammatory senescence-associated secretome, which perpetuates senescence and age-related pathologies through autocrine and paracrine signalling. Here, we show that T_EMRA_ cells from individuals with T2D express higher levels of all three TGFβ receptors together with increased amounts of TGFβ1i1. TGFβ1i1 functions as a transcription cofactor that along with Smad7 regulates p21 expression (69). Notably, glucose and metabolic dysregulation may influence TGFβ signalling and receptor expression (70). Therefore, the TGFβ-associated effects observed here should be considered within the context of an inflammatory and metabolically dysregulated T2D environment. However, we show here that TGFβ1 is a potential contributor of the altered T_EMRA_ phenotype observed in T2D, particularly through the internalisation of NK receptors. This aligns with previous findings demonstrating that TGFβ signalling in both mouse and human CD8^+^ T cells downregulated KLRG1 expression (71). Additionally, TGFβ may contribute to the observed oligoclonality in T_EMRA_ cells, as it promotes IL-7-dependent survival of low affinity T cells (72). Together, these findings reinforce the link between TGFβ-driven inflammation, metabolic dysregulation and premature senescence-like features in T2D-associated unhealthy immune ageing.

TGFβ not only shapes the phenotypic and functional properties of CD8^+^ T_EMRA_ cells but also modulates their tissue distribution by influencing adhesion molecule expression. In human plaque samples, CD8^+^ T_EMRA_ cells were represented at a higher frequency than in paired blood, while T2D-derived bulk CD8^+^ T cells showed greater splenic accumulation in the xenogeneic imaging model. These observations are consistent with altered tissue distribution. This may be driven by TGFβ-induced upregulation of CD103, a heterodimeric transmembrane complex that binds to E-cadherin facilitating tissue retention (73). Interestingly, E-cadherin also serves as a ligand for KLRG1, creating a competitive interaction between these two molecules. While KLRG1 engagement inhibits effector T cell function, CD103 binding to E-cadherin enhances cell-cell interactions (74). This acquisition of CD103 gives the T2D-associated T_EMRA_ population features reminiscent of tissue-resident-memory cells. Classical T_RM_ cells are generated from KLRG1^lo^ memory precursors, which are either KLRG1^-^ effectors or KLRG1^+^ effectors that have downregulated the receptor (ExKLRG1) and typically retain high cytotoxic and proliferative capacity (75). In contrast, the T2D-associated T_EMRA_ phenotype differs from this classical lineage by displaying reduced TCR-stimulated degranulation, suggesting an alternative differentiation state shaped by the metabolic disease environment. We propose that TGFβ may contribute to this phenotype by simultaneously inducing CD103 expression while downregulating KLRG1, a process documented to be dynamically regulated by intrinsic and extrinsic signals (75). Collectively, these findings underscore how a T2D-related inflammatory and metabolic environment promote a dysregulated, tissue-associated T_EMRA_ phenotype.

In addition to their altered tissue distribution and adhesion-associated features, CD8^+^ T_EMRA_ cells in T2D-associated unhealthy immune ageing exhibited a loss of cytotoxic potential, further compromising their ability to mount effective immune responses. TGFβ has been shown to suppress NK cell function, while knockout of the common signalling mediator SMAD4 enhances NK cell cytotoxicity (76). Consistent with the phenotypic data, CD8^+^ T_EMRA_ cells in T2D showed reduced Rab11a expression and altered receptor localisation, with TGFβ recapitulating part of this phenotype in vitro, suggesting that inflammatory signals may disrupt endocytic pathways through cytoskeletal remodelling. However, transcript-level changes alone do not fully establish the underlying trafficking mechanism and TGFβ-mediated downregulation of rab11A is unlikely to account for the entire reduction observed in T2D cells. High glucose and other metabolic factors present in T2D may also influence endosomal, lysosomal, and autophagy-associated pathways, potentially contributing to altered receptor trafficking dynamics. Efficient cytotoxicity requires co-ordinated trafficking and exocytosis of lytic granules, processes regulated by vesicular transport proteins such as the small GTPase rab27, which controls granule docking at the immunological synapse prior to degranulation (77). TGFβ signalling can downregulate the expression of rab family proteins, including rab27 via SMAD-dependent transcriptional suppression. Studies in breast cancer models have shown that TGFβ can reduce rab27 mRNA and protein levels through SMAD3-dependent mechanisms (78). In this disease-associated context, TGFβ could contribute to reduced rab27a expression and altered granule trafficking. TGFβ-mediated signalling could contribute to defective degranulation and diminished cytotoxic responses in CD8^+^ TEMRA cells.

Moreover, defects in receptor recycling and vesicular trafficking may also reflect the altered metabolic environment associated with T2D. As a metabolic disease characterised by dysregulated energy homeostasis, T2D may limit the energetic resources required for cytoskeletal rearrangement, vesicle transport, and membrane recycling (79). These pathways are tightly regulated by rab GTPases, which cycle between GTP-bound active and GDP-bound inactive states to coordinate vesicle docking, trafficking, and exocytosis (80). Impaired energy availability could therefore compromise GTP loading and rab activity, potentially disrupting receptor recycling and lytic granule exocytosis in CD8^+^ T_EMRA_ cells in T2D-associated unhealthy immune ageing.

Within T2D-associated unhealthy immune ageing, the disruption to cytotoxic function contributes to the accumulation of dysfunctional T_EMRA_ cells with altered capacity to provide effective local immune surveillance. In recent years, the experimental elimination of senescent cells has gained a lot of attention, as these studies show that the clearance of senescent cells increases healthy lifespan (81). Senescent cells can be recognised and eliminated by many different immune cell types including CD8^+^ T cells facilitated by their NK receptor expression (9). However, the appearance of a population of CD8^+^ T_EMRA_ cells which have reduced NK receptor expression and poor cytotoxic capability in unhealthy ageing may lead to incomplete elimination of senescent cells with age. It would be interesting to assess whether removing inflammation alone would be sufficient to restore CD8^+^ T_EMRA_ cells to their previous state or whether prolonged exposure to inflammation induces irreversible changes. However, even if inflammation removal allowed these cells to revert, achieving this clinically remains challenging, as the outcomes of anti-inflammatory therapies have been largely inconclusive. For instance, while anti-TNFα therapy has shown benefits in mouse models of insulin resistance and T2D (82), its therapeutic effects in humans have been limited (83, 84). Moreover, anti-TNFα treatment for rheumatoid arthritis is associated with an increased risk of serious infections (85). Furthermore, we show here that statins were associated with lower TGFβ levels in healthy older participants, but this association was not evident in the T2D group, suggesting that chronic inflammation alters TGFβ regulation in a way that resists conventional intervention. Consequently, a more targeted approach is needed to successfully manipulate CD8^+^ TEMRA cells in T2D-associated unhealthy immune ageing. Achieving this will require a deeper understanding of the molecular mechanisms driving their generation and persistence in this context.

## Acknowledgments

This work was supported by the British Heart Foundation (FS/15/69/32043, LAC), the Academy of Medical Sciences (SBF001\1013, ECC, SMH), Barts Charity (MGU0536, CGK, SMH and G-002143 JS, SMH), Diabetes UK (19/0006057, JB, SYAT, SMH) and the BBSRC (BB/X009610/1, VT, SMH). DA is supported by a Wellcome Trust Career Re-Entry Fellowship (221604/Z/20/Z DA) and Barts Charity (G-002145DA). DH is supported by the British Heart Foundation (BHF) Clinical Research Training Fellowship (FS/CRTF/20/24058) and FMB is supported by the BHF (CH/15/2/32064, RG/20/8/34995 and AA/18/5/34222). We thank the CRUK Flow Cytometry Core Service at Barts Cancer Institute (Core Award C16420/A18066). Additionally, this work was funded in part by the EPSRC programme for next generation molecular imaging and therapy with radionuclides (EP/S032789/1), the Wellcome/EPSRC Centre for Medical Engineering at King’s College London (WT 203148/Z/16/Z), a Wellcome Trust Multiuser Equipment Grant (212885/Z/18/Z). The nanoPET/CT scanner at KCL was funded by an equipment grant from the Wellcome Trust (WT 084052/Z/07/Z). Finally, we thank the research participants for their commitment to this study and their generous donation of blood samples.

## Author Contributions

CKG, LAC, SMH wrote the manuscript. CKG, LAC, JS, KL, ECC, DT, VSKT, IN, NER, BC, JB, MPDC ACM, DA, GPK, TTP, KS, RTMdR, SYAT, SMH designed and performed the experiments, as well as analysing the data and reviewed the manuscript. DH, FMB, CS, AW, GH, SF provided samples and reviewed the manuscript.

## Data availability statement

The data that support the findings of this study are available on request from the corresponding author. The data are not publicly available due to privacy or ethical restrictions.

## Supplementary Figure Legends

|  |  |  |  |  |  |  |
| --- | --- | --- | --- | --- | --- | --- |
| BMI range |  | 19-48 | 14-65 | 19-52 | 16-49 |  |
| Average BMI | | 28 ($\pm 6$) | 30 ($\pm 5$) | 30 ($\pm 7$) | 30 ($\pm 9$) | |
| Length of time with T2D (years) | | | 22 ($\pm 9$) | | | |
| CMV positivity |  | 61% | 66% |  |  |  |
| Ethnicity | White | 62% | 52% | 77% | 64% |  |
|  | Black, African, Caribbean or Black British | 8% | 14% | 11% | 12% |  |
|  | Asian or Asian British | 30% | 34% | 14% | 24% |  |
| Number of medications | 0 | 24% | 3% | 19% | 6% |  |
|  | 1-2 | 34% | 7% | 11% | 6% |  |
|  | 3-6 | 37% | 45% | 36% | 47% |  |
|  | 7+ | 5% | 45% | 33% | 41% |  |
| Type of medications | Diabetic | 0% | 91% | 0% | 94% |  |
|  | Cardiovascular | 45% | 79% | 78% | 100% |  |
|  | Statins | 41% | 83% | 31% | 56% |  |
|  | Gastrointestinal | 18% | 29% | 28% | 19% |  |
|  | Respiratory | 11% | 14% | 14% | 6% |  |
|  | Pain management | 12% | 17% | 9% | 8% |  |
|  | Other | 41% | 55% | 22% | 31% |  |

**Supplementary Figure 1.**
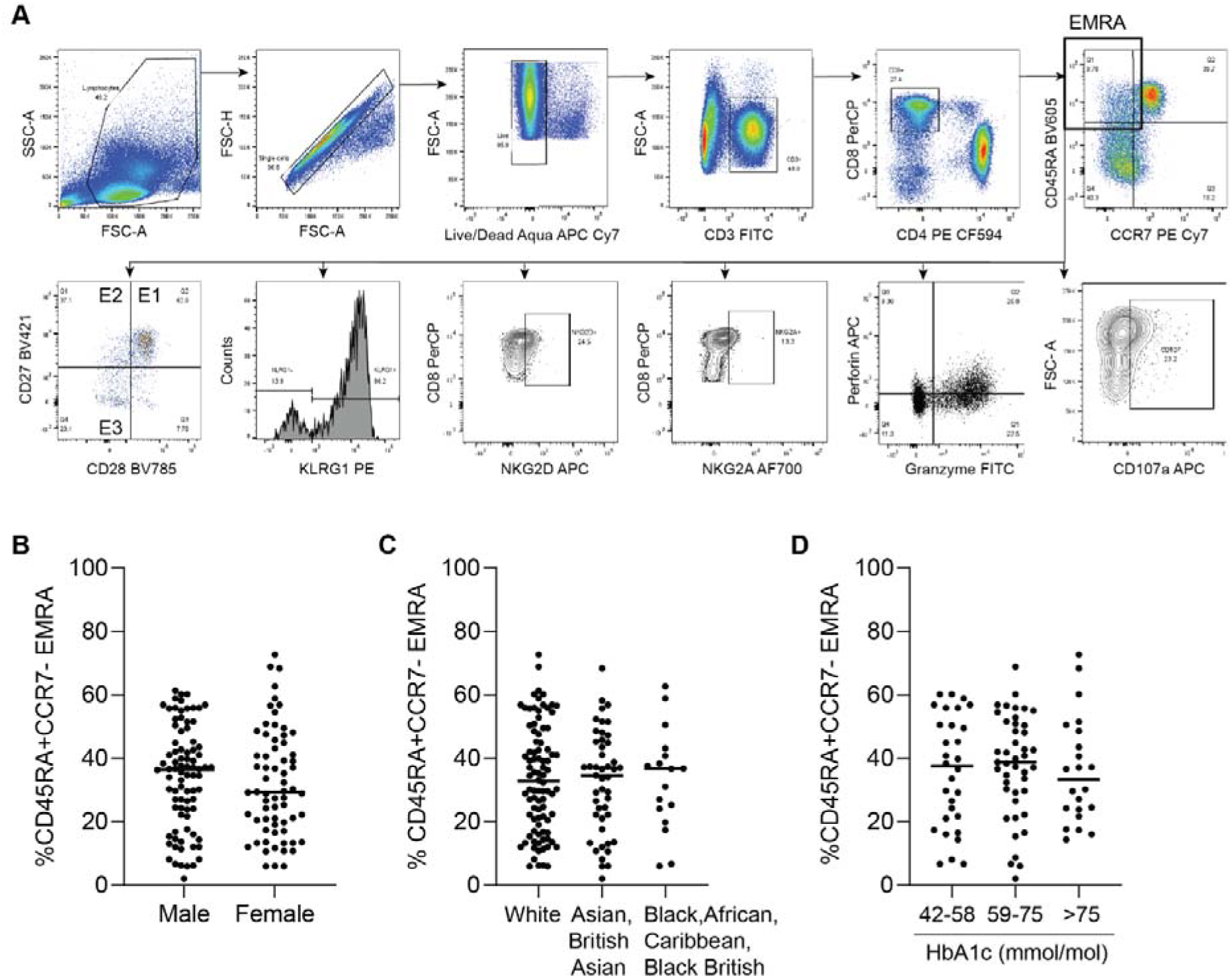
Phenotypic characterisation of CD8^+^ T cell subsets in T2D-associated unhealthy immune ageing. (A) Dot plots showing gating strategy. (B) CD45RA/CCR7 CD8^+^ T cells stratified according to gender and (C) ethnicity in the healthy and T2D cohorts. (D) CD45RA/CCR7 CD8^+^ T cells in the T2D cohort stratified by HbA1c (42-58, 59-75 and >75 mmol/mol).

**Supplementary Figure 2.**
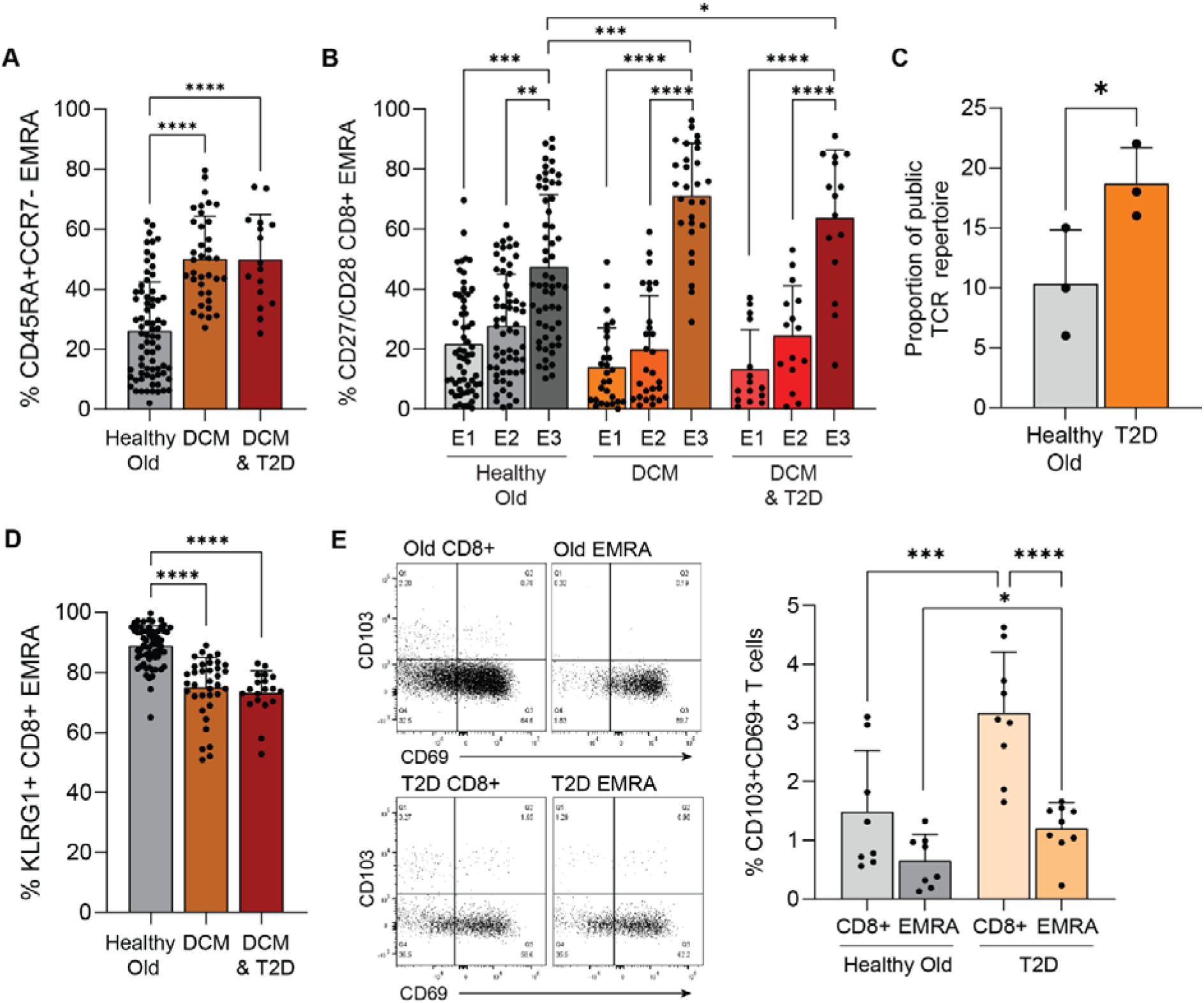
Characterisation of CD8^+^ T cell in dilated cardiomyopathy. (A) Graph showing CD45RA/CCR7 defined CD8^+^ T_EMRA_ cell populations and (B) CD27/CD28 expression in the CD45RA^+^CCR7^-^ T_EMRA_ population in individuals with dilated cardiomyopathy (DCM) with and without T2D compared to those without disease. (C) The proportion of the TCR repertoire with public sequences in people with and without T2D. (D) The proportion of T_EMRA_ cells that express KLRG1 in individuals with dilated cardiomyopathy with and without T2D compared to those without disease. (E) Flow cytometry plots and graph showing the proportion of CD103/CD69 defined T_RM_ cells in the total CD8^+^ population compared to the T subset. Data was analysed using ANOVA followed by Tukey multi-comparison test or a Mann-Whitney U test and expressed as mean ± SD. *p<0.05, **p<0.01, ****p<0.0001.

**Supplementary Figure 3.**
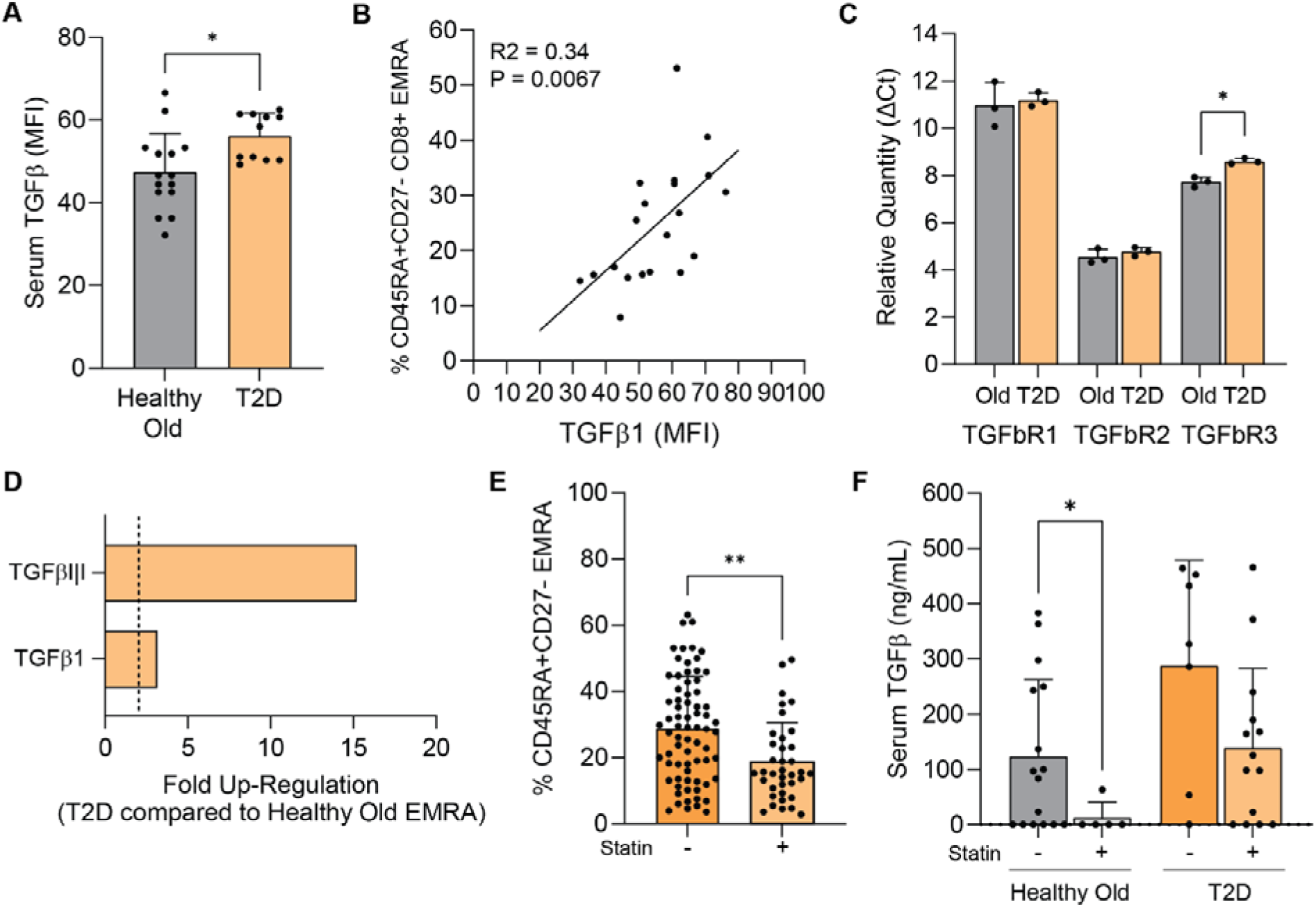
Association of TGFβ with CD8^+^ T_EMRA_ cells in T2D-associated unhealthy immune ageing. (A) Serum TGFβ levels measured by Legend plex in individuals with and without T2D. (B) Correlation between serum TGFβ levels and the proportion of CD45RA/CD27 defined CD8^+^ T_EMRA_ cells. A line of best fit was generated using simple linear regression analysis. The coefficient of determination (R^2^ = 0.34, p = 0.0067) is shown to indicate goodness of fit. (C) Relative gene expression of TGFβ receptors using RT-PCR in T_EMRA_ cells from individuals with and without T2D. (D) Graph showing the upregulation of TGFβ and TGFβI|I genes in T_EMRA_ cells using an RT^2^ profiler PCR array. Genes with a fold change ≥ 2 were considered significant. Data compares individuals with and without T2D, n = 3 for each group. (E) Graph showing the proportion of CD45RA/CD27 defined T_EMRA_ cells in individuals taking statins compared to those who do not. (F) Serum TGFβ levels measured by Legend plex split into individuals with and without T2D and whether they take statins.

**Supplementary Figure 4.**
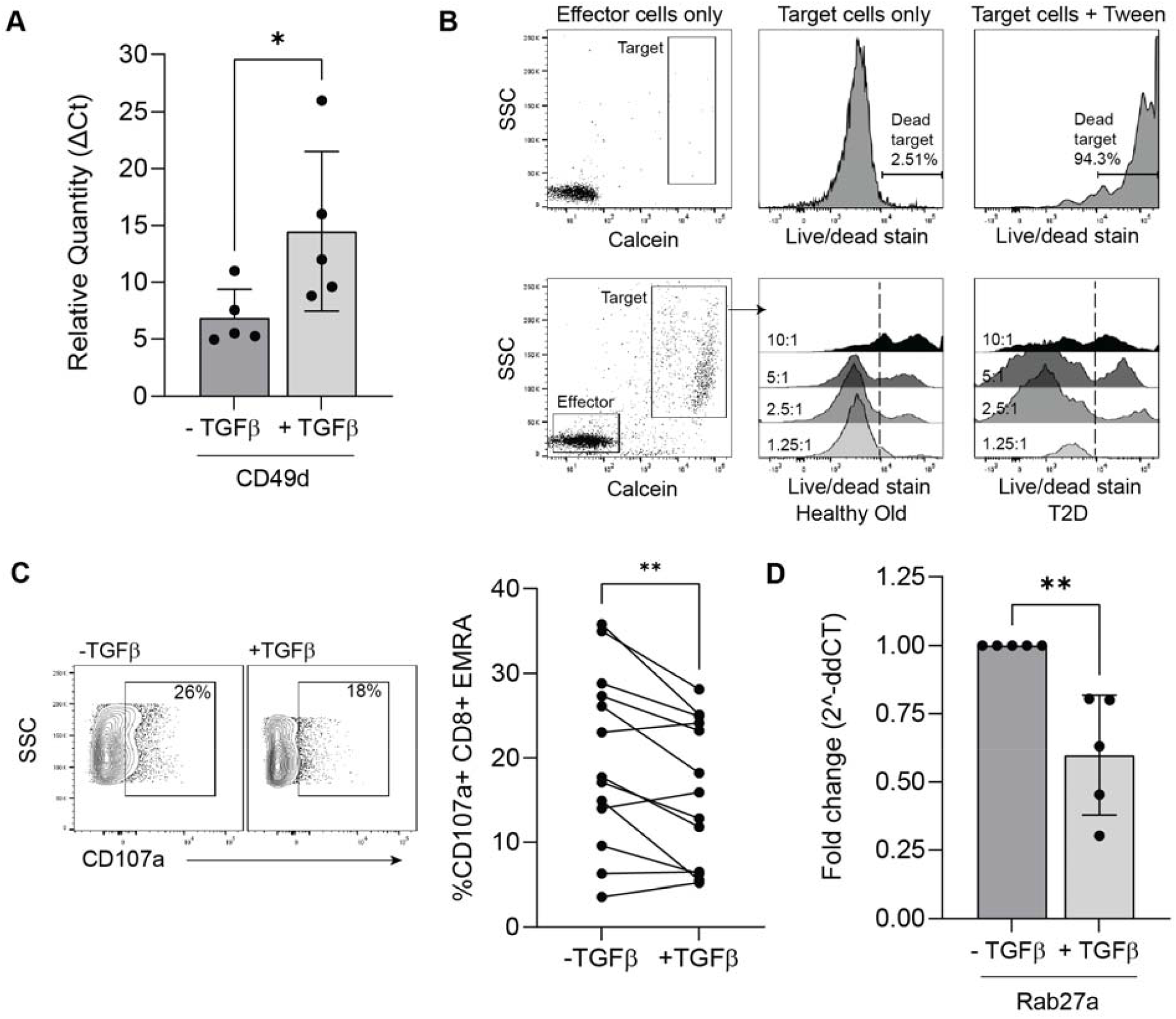
(A) Relative mRNA expression of CD49d in CD8^+^ T cells incubated with and without TGFβ1 for 5 days. Expression was normalised to βActin and calculated using the ΔCt method. (B) The upper row shows control conditions for the cytotoxicity assay: effector cells only, target cells only, and target cells treated with 0.1% Tween-20. The bottom row displays a representative anti-CD3-stimulated bulk CD8^+^ T-cell cytotoxicity assay, showing target cell death at effector to target (E:T) ratios of 1.25:1, 2.5:1, 5:1, and 10:1. (C) Flow cytometry plot and graph showing anti-CD3-stimulated CD107a externalisation in CD45RA/CCR7 defined T_EMRA_ cells with and without 10 ng/mL TGFβ1. (D) Relative mRNA expression of rab27a in CD8^+^ T cells incubated with and without TGFβ1 for 5 days. All data was analysed using a Mann-Whitney U test and is expressed as mean ± SD. *p<0.05 and **p<0.01.

**Supplementary Figure 5.**
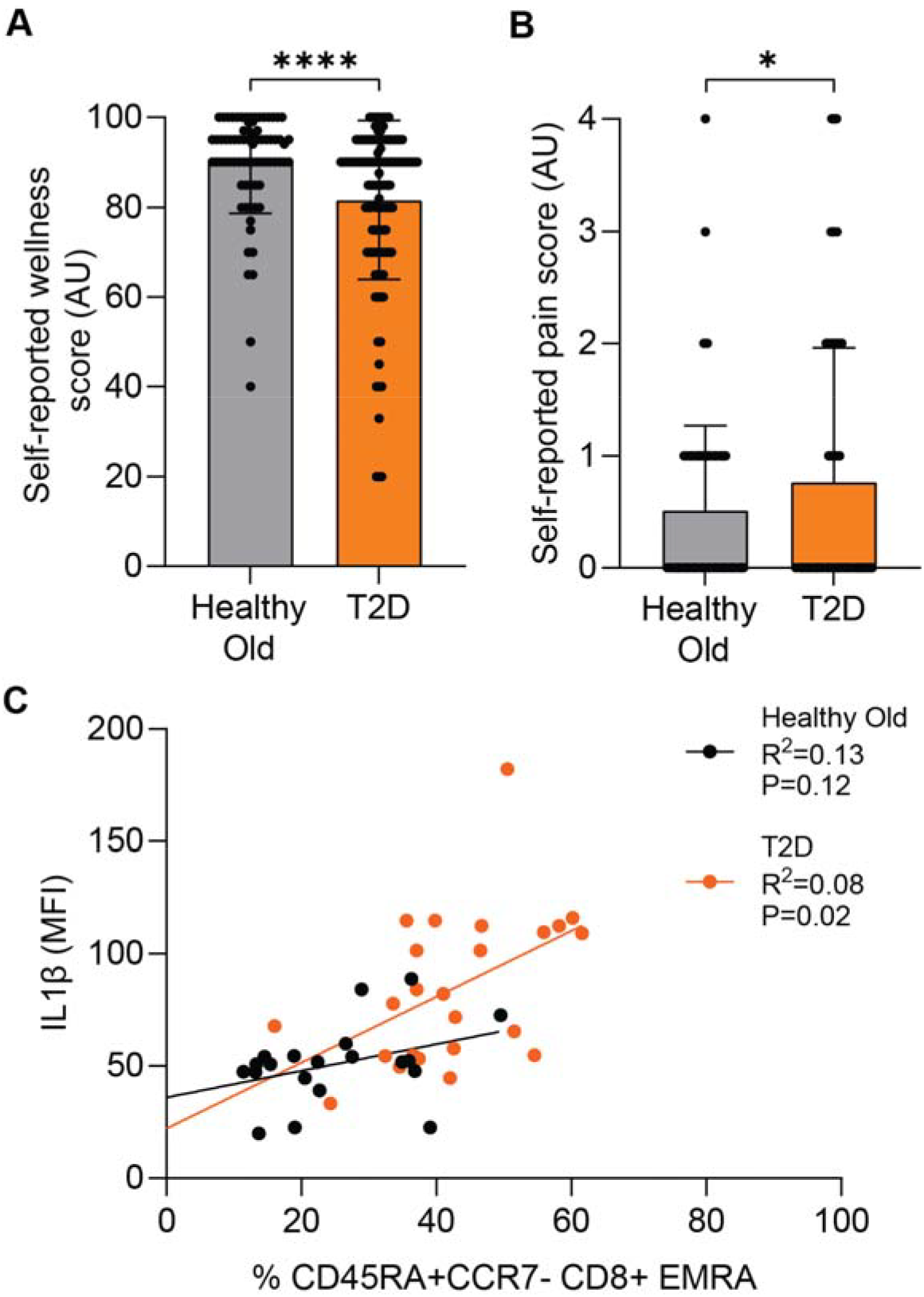
Reduced self-reported wellbeing in individuals with T2D. (A) EQ-5D visual analogue scale (VAS) score assessing perceived health status on the day of assessment. Scores range from 0 to 100, where 100 indicates the best health imaginable and 0 indicates the worst. (B) Self-reported pain and discomfort score derived from the EQ-5D descriptive system: 0, no pain or discomfort; 1, slight pain or discomfort; 2, moderate pain or discomfort; 3, severe pain or discomfort; 4, extreme pain or discomfort. (C) Correlation between serum IL-1β concentrations and the proportion of CD45RA/CD27-defined CD8^+^ TEMRA cells in individuals with and without T2D. Lines represent the best fit from simple linear regression analysis and the coefficient of determination and P values are shown to indicate the strength and statistical significance of the association.

